# Rapid, replicated field tissue collection for population-scale plant omics

**DOI:** 10.64898/2026.11.28.769293

**Authors:** Jonathan Turkus, James C. Schnable

## Abstract

Measuring gene expression and other molecular phenotypes across populations can improve trait prediction and uncover trait-controlling genes missed by DNA markers. Generating such datasets in the field requires collecting and immediately freezing hundreds to thousands of tissue samples within a short window before diurnal shifts confound biological signal. Here we describe an open-source field tissue collection kit combining a triple leaf punch, hot-swappable tube trays, a sealed cryogen carrier, and a wearable or wheeled carrier box. Across nine field campaigns in three states, the kit enabled rapid, triplicate sampling of up to 1,050 plots in 75 minutes. In a 966-plot sorghum experiment sampled in 92 minutes, collection time explained 1.1% of expression variance for the median gene, versus a projected 15.4% over six hours. Dry ice yielded transcriptomes indistinguishable from liquid nitrogen flash freezing, and between-plot carryover was below one read in 20,000 without punch sterilization. Design files for 3D-printed parts, written fabrication and field protocols, assembly videos, and translations of this manuscript in 23 languages are openly available.

## 1 Introduction

Molecular phenotypes mediate genetic variation and whole-plant traits [9, 26, 33]. Population-scale measurements of metabolite abundance and/or gene expression datasets have been shown to, in some cases, improve trait prediction relative to methods based solely on DNA markers [4, 10, 20, 33]. In addition, measuring gene expression across populations has been used to identify trait-associated genes missed by DNA markers [23, 24, 40], potentially by integrating weak *trans*-acting variants [24] or resolving multiallelic *cis*-regulatory variant loci that would otherwise be collapsed by individual biallelic markers [14].

Sequencing costs have historically represented the primary barrier to measuring gene expression levels across large populations. Costs of both library preparation and sequencing have fallen to the point that measuring gene expression across a 1,000-genotype population can plausibly be accomplished within the budget of an individual research lab [2, 21, 29]. However, datasets of this scale remain scarce, in part because of the greater logistical challenge that collecting samples suitable for quantifying labile molecular phenotypes represents relative to collecting samples suitable for genotyping DNA differences.

Excising a leaf sample induces transcriptional and metabolic responses within minutes [8, 16, 38], so measuring molecular phenotypes as they exist in the field requires immediate freezing to mitigate wounding- and water-loss-induced responses. Once frozen, samples must be maintained in a frozen state until extraction to avoid degradation of transcripts and biochemical changes resulting from lysed cells, requiring a reliable cold chain from the field (or greenhouse) to the molecular biology lab. In addition, gene expression and metabolite abundance exhibit strong environmental, batch, and diurnal effects that can obscure trait associations [17, 19, 22, 36]. Transcript levels shift substantially across plants collected at different times of day in the same experiment [17, 22]. This means population-scale expression datasets are not built incrementally from samples collected in different grow-outs or as part of different collection campaigns, in contrast to DNA marker panels where joint analyses of data generated in different research groups can produce datasets for increasingly larger populations [3, 6, 15]. This also means tissue samples that fail or are lost during extraction or quantification cannot be informatively recollected to fill gaps in a dataset, making collecting backup samples as part of collection campaigns for labile molecular phenotypes critical and further slowing collection efforts for these campaigns.

Here we describe the design, assembly, and evaluation of a sample collection kit that can be constructed from a mixture of widely available commercial components and 3D-printed parts that can be employed to collect and flash freeze large numbers of plant samples in triplicate, providing built-in backups against lost samples or failed quantification and/or the ability to profile metabolism and gene expression from inherently paired samples. Analysis of transcriptomic samples collected using the system confirms a lack of significant carryover between sequential plots and demonstrates that gene expression measured from samples collected onto dry ice using this kit is essentially indistinguishable from those flash frozen with liquid nitrogen, reducing a significant labor and logistical challenge to quantifying gene expression from large experiments at remote field sites. As part of this manuscript, we have developed and released written fabrication and collection protocols, video walkthroughs, and translations of this manuscript in a total of 23 languages representing a cumulative 3.87 billion native speakers around the globe (Supplementary Table S3).

## 2 Results

### 2.1 A system for collecting replicated tissue samples at field scale

The sampling kit integrates four components: a triple leaf punch, field-swappable tube trays, a cryogen carrier, and a tote that allows the tube tray and cryogen carrier to be easily transported together (Figure 1). The triple leaf punch holds three tubes simultaneously and deposits one leaf disk into each with every squeeze, producing three matched samples from the same leaf. The tube trays are pre-loaded with labeled tubes in the planned order of collection. The tray is engineered to grip the tubes tightly enough that they will not fall out during transport or collection, even if fully inverted, but loosely enough and specifically positioned such that all three tubes can be loaded into the triple punch in a single motion prior to sample collection. After collection, all three tubes can be removed from the triple punch in another single motion, capped, and then deposited into the cryogen carrier via a trap door that keeps the tubes and cryogen from escaping the carrier, even if the kit is dropped or the collector trips in the field, while also limiting the flow of warm air into the sample/cryogen compartment. The tote used to transport the cryogen carrier and tube trays can be carried on a strap worn around the neck of a single sample collector or pulled on a collapsible cart (more comfortable for long campaigns or two-person teams). Design files and assembly protocols are provided for all of these components (see Data availability statement).

**Figure 1:**
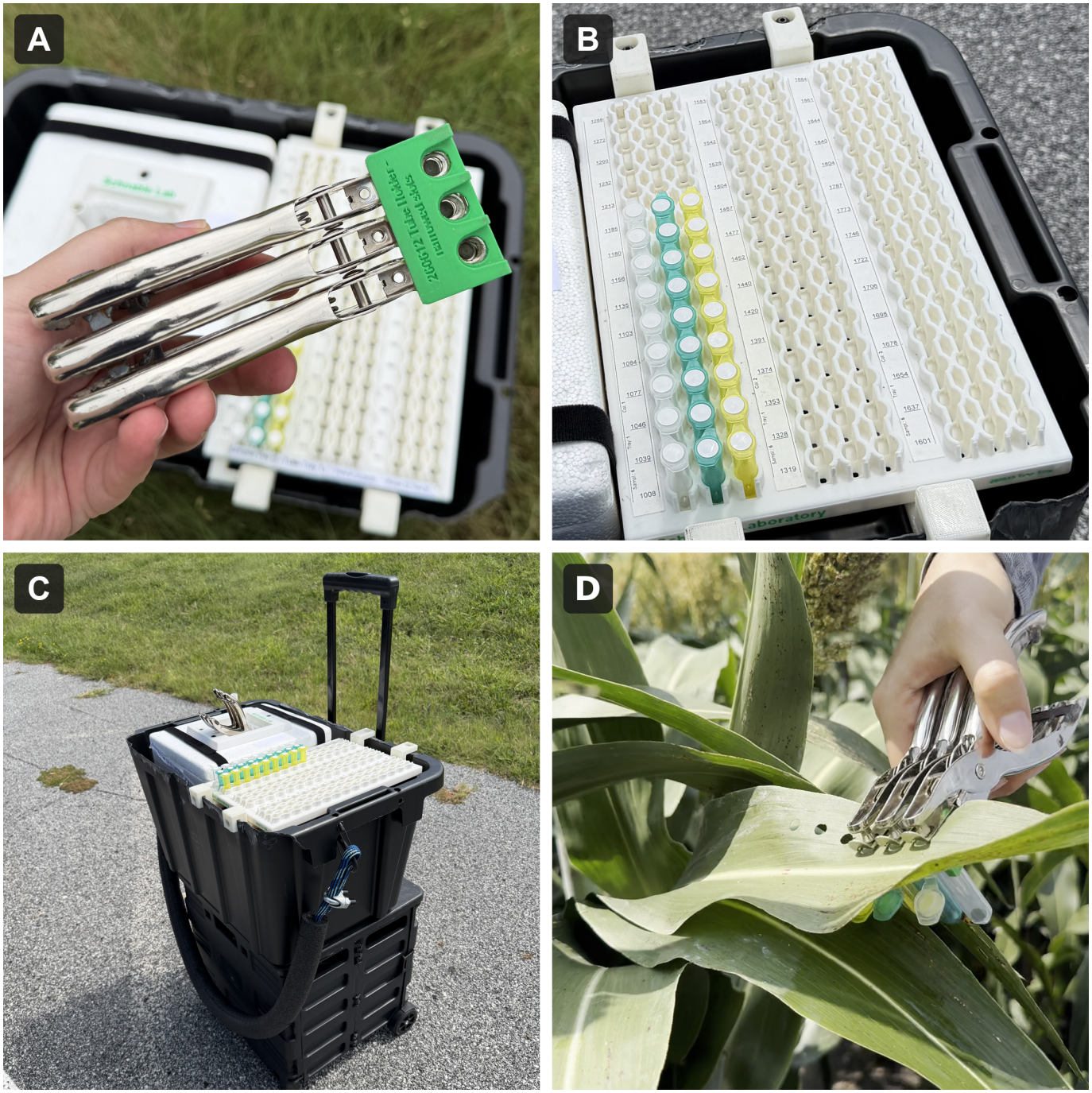
The rapid field tissue collection kit. **(A)** The triple leaf punch: three single-hole punch bodies joined with the 3D-printed triple punch adapter, held above an open carrier box containing the cryogen carrier and a loaded tube tray. **(B)** A 3D-printed tube tray seated on the tray holder along the open side of the carrier box, holding pre-labeled tubes in collection order. **(C)** Complete kit ready for collection with cryogen carrier, tube trays, carry strap, and collapsible cart. **(D)** The loaded triple punch collecting leaf disks from sorghum in the field.

Across nine field campaigns conducted over a five-year period where detailed timing records were kept, individual samplers frequently sustained collection rates of >60 plots/hour (Supplementary Figure S1). Two-person teams using one kit sustained collection rates of >80 plots/hour per team. Campaign-level rates were somewhat lower because total duration was constrained by the slowest sampler, staggered starts, and uneven numbers of plots collected by different samplers (Table 1). Overall, individual samplers working with neck straps are an optimal solution when collection rate is limited by the number of people available to participate in a campaign, while two people working together from a single kit provide benefits when fewer collection kits are available than people.

**Table 1:** Example sampling campaigns and collection speeds. Rates are plots per hour of active sampling / plots per hour from the start to the end of the campaign. NE, Nebraska; NC, North Carolina; SD, South Dakota.

| Year | Crop | Plots | Kits | People | Minutes | Plots person<sup>-1</sup> h<sup>-1</sup> | Plots kit<sup>-1</sup> h<sup>-1</sup> |
|---|---|---|---|---|---|---|---|
| 2021 | Maize | 840 | 11 | 12 | 112 | 59 / 38 | 62 / 41 |
| 2021 | Sorghum | 966 | 11 | 14 | 92 | 65 / 45 | 65 / 57 |
| 2022 | Maize | 385 | 7 | 7 | 64 | 72 / 52 | 72 / 52 |
| 2023 | Maize-NE | 300 | 4 | 4 | 96 | 59 / 47 | 59 / 47 |
| 2023 | Maize-NC | 250 | 3 | 3 | 67 | 78 / 75 | 78 / 75 |
| 2023 | Sorghum | 762 | 8 | 15 | 117 | 36 / 26 | 59 / 49 |
| 2026 | Maize | 814 | 8 | 16<sup>a</sup> | 91 | 40 / 34 | 80 / 67 |
| 2026 | Sorghum-NE | 1,050 | 11 | 22<sup>a</sup> | 75 | 44 / 38 | 88 / 76 |
| 2026 | Sorghum-SD | 1,050 | 11 | 11 | 126 | 52 / 45 | 52 / 45 |

<sup>a</sup> Two-person teams.

### 2.2 Impact of time of collection on gene expression

The use of preloaded and labeled tubes requires that the order of sample collection for each participant be determined in advance. By recording individual participants’ start and stop times as well as, optionally, the time at which each empty tube tray is swapped out for a filled one, the time of collection for each individual sample can be inferred to minute-level resolution. The high-resolution inferred timing of each sample makes it possible to observe how rapidly gene expression can change over time, even within short and defined sampling windows. A 2021 sampling campaign [31] used the collection kits to collect samples from more than 900 sorghum plots in approximately one and a half hours (Table 1). Over the 92-minute span from the collection of the first to the last sample, the expression of *SbLHY* (Sobic.007G047400), a component of the sorghum circadian clock, declined from approximately 2,650 to approximately 1,100 TPM, with collection time inferred from start/stop/swap times explaining 47% of variance in the expression of this gene (Figure 2A).

**Figure 2:**
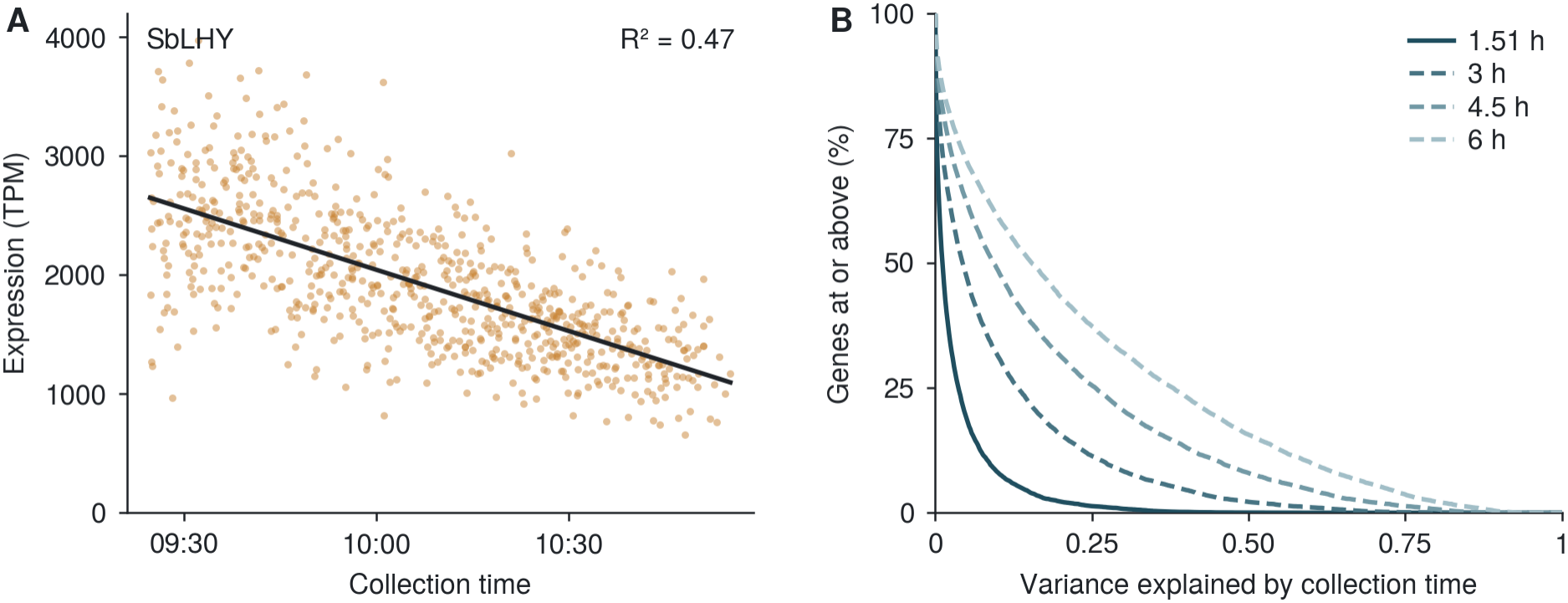
Impact of collection time on gene expression in the 2021 Nebraska sorghum field. **(A)** Expression of *SbLHY* (Sobic.007G047400) across 735 RNA-seq libraries collected between 09:24 and 10:56. **(B)** Cumulative distribution of the proportion of expression variance explained by collection time across 9,822 expressed genes (mean *≥* 10 TPM). The solid curve shows the observed 92-minute window; dashed curves show linear projections for 3, 4.5, and 6 h windows.

Across the set of 9,822 sorghum genes with mean expression *≥* 10 TPM in this dataset, 62.3% exhibited a statistically significant correlation between expression and inferred sample collection time (FDR <0.05) with a median rate of change across all 9,822 genes of approximately 10% of mean expression per hour. Because of the rapid collection window, the median proportion of total variance in gene expression attributable to time of collection was only 1.1%; however, projecting observed changes to longer time intervals suggested that over a 6-hour collection window time of collection would be responsible for at least 15% of observed variance in gene expression. Estimates using gene expression data from a separate maize sample collection campaign gave similar results (Supplementary Figure S2).

### 2.3 Substitution of dry ice for liquid nitrogen

The typical cryogen used to flash freeze plant tissue samples intended for gene expression or metabolite assays is liquid nitrogen. Dry ice is both much more widely available away from university campuses and presents fewer handling hazards than liquid nitrogen. However, both the difference in temperature (*−*78.5 °C for dry ice vs. *−*196 ^*°*^C for liquid nitrogen) and the fact that samples can be completely immersed in liquid nitrogen vs. reduced physical contact with solid dry ice indicate that the process of freezing is likely to be more prolonged when using dry ice as a cryogen, potentially providing additional time for wounding-related transcriptional and metabolic responses and/or inducing chilling-related responses. Some studies omit freezing entirely and substitute immersion of the sample in RNA preservative solutions that use high concentrations of salt to inactivate RNases.

In a pilot study conducted using replicated samples collected from the same plant at the same time using either the collection kit with liquid nitrogen in the cryogen carrier, a duplicate kit with dry ice in the cryogen carrier, or an empty cryogen carrier but with a commercially available RNA preservation solution pre-loaded in the sample collection tubes, samples preserved in a commercial RNA preservative without the use of a cryogen exhibited sharply distinct gene expression profiles from samples collected using either dry ice or liquid nitrogen, as well as substantially higher variance between replicated samples collected from the same plant at the same time (Supplementary Figure S3A). Differential gene expression analysis identified more than 7,000 genes (one third of those tested) as either up- or downregulated in the RNA preservative collected samples relative to either the liquid nitrogen or dry ice samples (Supplementary Figure S3B).

An expanded test examined samples collected from 25 maize hybrid plots grown as part of the Lincoln, Nebraska field site of the Genomes to Fields experiment in 2023 [7]. As above, samples were collected from the same plants using two collection kits, one with a cryogen carrier carrying liquid nitrogen and one with a cryogen carrier carrying dry ice. RNA concentration and RIN values were indistinguishable between cryogens (Supplementary Table S1). Gene expression data from paired samples collected using different cryogens were tightly clustered (Figure 3A), with cryogen explaining <1% of total variance in gene expression (Figure 3B). The median correlation of gene expression between samples collected from the same plot with different cryogens was *R*^2^ = 0.97 while the median correlation in gene expression between samples collected from different plots representing different maize hybrids was *R*^2^ = 0.83 (Figure 3C).

**Figure 3:**
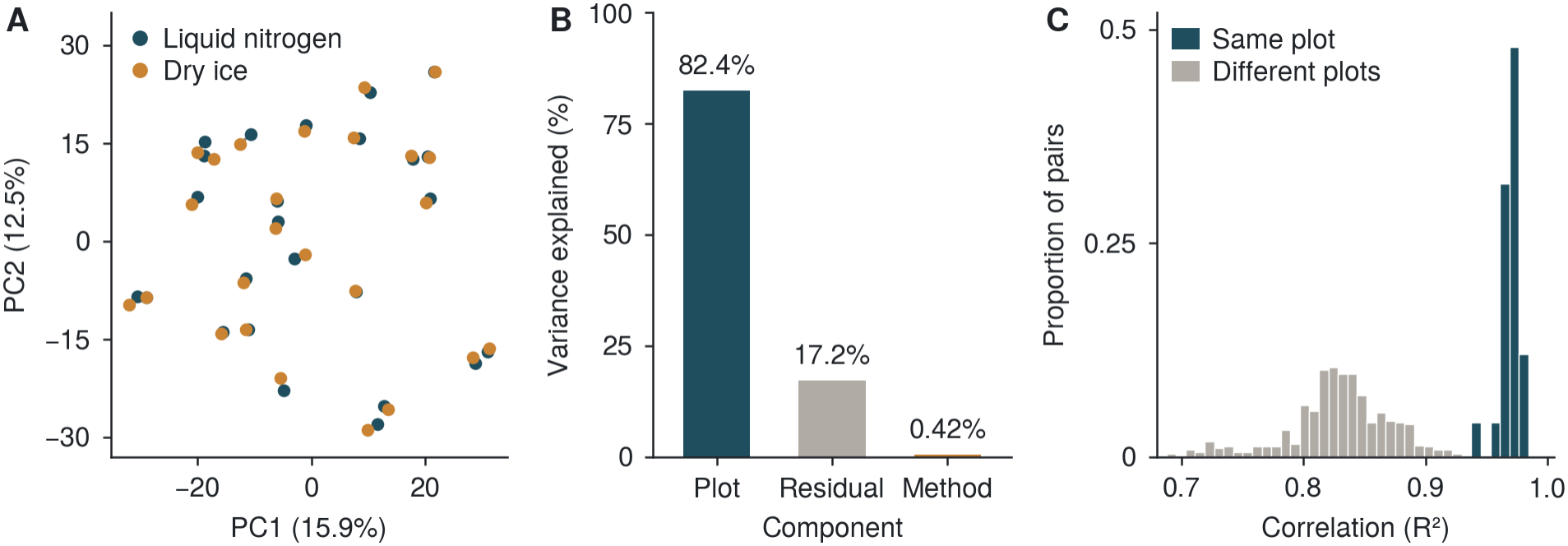
Dry ice and liquid nitrogen yield equivalent maize leaf transcriptomes. **(A)** PCA of the 500 most variable genes across 50 RNA-seq libraries collected on liquid nitrogen or dry ice from 25 hybrid maize plots; libraries from the same plot fall nearly on top of each other. **(B)** Partitioning of expression variance into plot, collection method, and residual components. **(C)** Expression *R*^2^ between liquid nitrogen and dry ice libraries from the same plot versus all mismatched plot pairs.

### 2.4 Quantifying carryover between sequentially collected field samples

A key step that can slow sample collection in the field is sterilization of tissue collection equipment between each sample. The protocol proposed and employed for these leaf collection kits omits a sterilization step and replaces it with a quick application of the triple punch to a plant from the same plot as the plant being sampled (or the same plant) prior to loading the sample collection tubes into the punch (Supplementary Protocol S2). Gene expression data generated from seven pairs of sequentially collected plots representing different maize genotypes, each of which had been previously genotyped via whole-genome resequencing, were evaluated to quantify how much carryover occurred using this protocol. Examining single nucleotide polymorphisms where sequentially sampled maize genotypes carry different alleles provides a mechanism to quantify cross contamination. At the subset of these sites located in expressed genes, 0.086% of aligned reads from the second plot in sequentially sampled pairs carried alleles specific to the first plot of the same pair, while 0.081% of aligned reads from the first plot carried alleles specific to the second plot. Because tissue or RNA from a later sample cannot enter the tube of an earlier collected sample via cross contamination, the 0.081% likely represents contamination introduced during sample processing, extraction, and library construction and/or misassignment of reads between multiplexed libraries during sequencing. The difference between these two proportions, about 0.005% of aligned reads at informative sites (corresponding to one read per 20,000) is our estimate of the prevalence of carryover from field collection using the sample collection kit and protocol described here.

## 3 Discussion

Large collaborative multi-environment field experiments such as Genomes to Fields [1, 7], SoyNAM [12], DROPS [34], and CUBIC [28] have created datasets at a scale never before available in the public sector and have enabled researchers to make significant advances in modeling and studying how genetic and environmental factors govern plant phenotypes and crop performance, both via analyses conducted by members of the collaborations and via data reuse and open prediction contests [7, 27]. While measuring transcript abundance at individual field sites has shown promise in identifying genes responsible for controlling variation in plant phenotypes as well as improving phenotype prediction [4, 10, 20, 23, 24, 40], dissecting genotype-by-environment interactions requires profiling these traits across multiple environments, and datasets comparable in scale to those described above do not yet exist for molecular phenotypes.

Tools that address the logistical challenges of collecting tissue samples suitable for molecular phenotyping can help close this gap by lowering barriers to participation and increasing the number of research groups able to undertake collections and studies. Increasing the speed of sample collection improves the quality and usefulness of molecular trait data by limiting variance introduced by diurnal cycling and changes in temperature and sunlight throughout the day (Figure 2). Faster sampling also puts measuring molecular phenotypes within the capabilities of smaller research teams: in North Carolina in 2023, it took only three people and three kits to sample 250 plots in 67 minutes (Table 1). Because liquid nitrogen can be difficult to obtain away from major research universities, being able to confidently substitute dry ice (widely available through consumer retail channels) lowers barriers at remote field sites. Much of the hands-on work of field crop trials worldwide is carried out by technicians, students, and agronomists working in languages other than English, so providing assembly instructions, standardized protocols, and this manuscript in 23 languages (Supplementary Table S3) lowers an additional training and adoption barrier.

While the initial motivation to collect samples in triplicate was to ensure backups for samples lost during processing, in our experience the third sample (second backup) is rarely, if ever, required for this purpose. These samples can be saved for long periods of time, as the samples from a 1,000-plot trial occupy only 12 L of *−*80 ^*°*^C freezer space, allowing a single upright *−*80 ^*°*^C freezer to archive approximately 30 site-years of tissue. Archiving these backup tissue samples across growing seasons creates biobanks that can be tapped into by future projects to study multi-year environmental responses employing future molecular assays.

## 4 Materials and Methods

### 4.1 Sampling kit fabrication and field use

The triple leaf punch joins three single-hole tissue punches (BioSpec, Cat. #709; silicone grips and center spring removed to reduce actuation force) aligned in a 3D-printed adapter and tack-welded with two mild steel strips. The 3D-printed tube tray mounts via hook-and-loop strips onto a 3D-printed tray holder bolted inside a modified polypropylene storage tote (interior *≈* 17 *×*11 *×*11 in). The cryogen carrier is built from an expanded-polystyrene shipping box (Uline, Cat. #S-18313) fitted with a CNC-routed HDPE trap door, 3D-printed faceplate and spacers, M5 hardware, a brass hinge, an extension spring, and hook-and-loop lid straps. Fabrication requires a 3D printer, CNC router, metal-cutting tool, MIG welder, drill press, and hot knife, and takes *≈* 2 person-hours per kit after part printing and cutting. Complete parts lists, safety notes, step-by-step fabrication instructions, and the field collection procedure are provided in Supplementary Protocols S1 and S2.

### 4.2 Field collections and RNA-seq

RNA-seq analyses used three Nebraska field experiments grown at the University of Nebraska– Lincoln Havelock Farm (40.86^*°*^ N, 96.60^*°*^ W): the 2021 Wisconsin Diversity maize panel [32] (planted May 7, collected July 8, 2021), the 2021 Sorghum BioDiversity panel [13, 31] (planted May 25, collected August 5, 2021), and 25 hybrids from the 2023 eastern Nebraska Genomes to Fields trial [7]. The 2021 maize and sorghum campaigns used the full kit with liquid nitrogen. In the 2023 trial, each of the 25 plots was sampled simultaneously with paired liquid nitrogen and dry ice kits, and a preservation pilot compared liquid nitrogen, dry ice, and RNAlater, three replicates each. RNA was extracted with the MagMAX Plant RNA Isolation Kit (ThermoFisher Scientific, A47157) and sequenced as paired-end (2 *×* 150 bp) Illumina RNA-seq libraries.

### 4.3 Gene expression quantification

Gene expression was quantified with kallisto v0.48 [5] against Phytozome v13 transcriptomes for maize B73_RefGen_v5 (Zm-B73-REFERENCE-NAM-5.0 v5.5) [18] and sorghum BTx623 (Sbicolor v5.1) [35], and summarized to gene-level TPM and counts via tximport [39] in R [37]. Collection-time analyses were restricted to genes averaging *≥* 10 TPM.

### 4.4 Cryogen, collection-time, and differential expression analyses

For the 25-plot liquid nitrogen versus dry ice comparison, genes with TPM *≥* 1 in at least 10 of the 50 libraries were retained and log_2_(TPM + 1)-transformed. PCA was performed on the 500 most variable genes (centered, unscaled), expression variance was partitioned across plot, cryogen, and residual components by method-of-moments ANOVA pooled across genes, and pairwise *R*^2^ values were computed across all 25 matched and 600 mismatched plot pairs. Differential gene expression in the preservation pilot was tested with DESeq2 [30].

Per-plot collection times in the 2021 maize and sorghum campaigns were inferred from planned plot order and recorded sampler start, tray-swap, and end times, assuming a constant sampling rate within each tray while allowing rates to vary between trays. Single-gene time trends were estimated by linear regression of expression (TPM) on inferred collection time. Variance partitioning and linear projections across 3, 4.5, and 6 h windows included sampler as a fixed effect, so that differences between samplers were not attributed to collection time.

### 4.5 Carryover analysis

Carryover was estimated from the first two plots collected by seven samplers in the 2021 Nebraska maize experiment whose genotypes were present in a published B73_RefGen_v5 whole-genome resequencing (WGS) marker set [15] and passed RNA-seq QC. Reads were aligned to B73_RefGen_v5 with bwa mem v0.7.17-r1188 (`-M`) [25], duplicates were marked with samtools v1.19.2, and allele counts were extracted with bcftools v1.19 `mpileup` (mapping quality *≥*20, base quality *≥*20) [11]. Sites were retained if covered by *≥*10 unique reads in both samples and if *≤* 1% of reads carried unexpected alleles, excluding loci prone to paramorphisms. To control for reference bias, carry-over rates were calculated separately depending on whether the first or second sample carried the reference allele, and compared against reciprocal rates and third-allele error rates (Supplementary Table S2).

## Data availability

Design files for the custom kit components and the fabrication and field collection protocols (Supplementary Protocols S1 and S2) in 23 languages, and translations of this manuscript into 22 languages, are available from figshare (https://doi.org/10.6084/m9.figshare.35094379; Supplementary Table S3). How-to videos for kit fabrication in the same 23 languages are available at https://www.youtube.com/watch?v=Ozwe9-aWtT0&list=PLGAixDAU2zuQ. A video demonstration of the field collection procedure is available at https://www.youtube.com/watch?v=ep0gtoMmKBw. FASTQ files for the 2021 Nebraska sorghum RNA-seq dataset are deposited in the European Nucleotide Archive under accession PRJEB83049 [31].

## Supplementary information

Supplementary Tables S1–S3, Supplementary Figures S1–S3, and Supplementary Protocols S1–S2 are provided in the accompanying Supplementary Information.

## Author contributions

J.T. designed and fabricated the original system and subsequent revisions, contributed to data collection, and designed the cryogen experiment. J.C.S. conducted the data analyses and drafted the paper. Both authors revised and approved the final manuscript.

## Acknowledgements

The authors thank Aleah Miller, Alexa Nolan, Alice Guo, Alliance Igiraneza, Amany Gomaa, Annika Waller, Anuradha Choudhary, Babou Clion Muhoza, Bo Peng, Bri Johnson, Brooke Bouwens, Chidanand Ullagaddi, Christopher Wang, Daniella Norah Tumusiine, Deniz Istipliler, Emily Francis, Ermias Assefa, Fangyi Li, Guangchao Sun, Hadiya Kounsar, Han Tran, Hannah Pil, Harshita Mangal, Hector Palala, Henrique Moura Dias, Hongyu Jin, Insoo Jeon, Isabel Sigmon, Jensina Davis, Joe Gage, Jordan Bares, Kangtoe Lee, Karla Montserrat Cuéllar Pérez, Katelyn Fritz, Khubi Basnet, Kyle Linders, Lina López-Corona, Mackenzie Zwiener, Makenzie Crum, Marcin Grzybowski, Matt Vredenburg, Meijie Luo, Michael Tross, Muragesh M. Hiremath, Muskan Jakhar, Nate Korth, Nikee Shrestha, Özgür Altundaş, Prajwal R. Shashishekhar, Prince Ngiruwonsanga, Ramesh Kanna Mathivanan, Ravi Mural, Ryleigh Kirby, Sachina B.K., Saet-Byul Kim, Sangsik Yun, Sawyer Johnson, Shalma Maman, Shiva Kumar Reddy Mudedla, Sophie Rohlfing, Srimathi Bala, Sumeet Mankar, Taryn Peffers, Tricia King, Ty Thomas, Vladimir Torres-Rodriguez, Waqar Ali, and Zhongjie Ji for contributing to the field sampling campaigns and providing feedback that improved the kit and protocol. We also thank Anna Podgórska-Grzybowska, Anuradha Choudhary, Bibash Rokaya, Fotima Turdalieva, Harshita Mangal, Henrique Moura Dias, Karla Montserrat Cuéllar Pérez, Marcin Grzybowski, Masako Obata, Muragesh M. Hiremath, Özgür Altundaş, Pritha Kundu, Sai Subhash Mahamkali VS, Shokhista Turdalieva, Surya Krishna Sakthivel, Waqar Ali, and Zhongjie Ji for providing narration.

## Ethics approval and consent to participate

Not applicable.

## Consent for publication

Not applicable.

## Use of artificial intelligence

AI models including, but not limited to, Gemini 4 Neon, Claude Opus 5.5, and ChatGPT Sol 6.1 were used to assist in data analysis, visualization, and video and textual editing. The authors take full responsibility for the integrity of all results presented in this work.

## References

[1] N. AlKhalifah, D. A. Campbell, C. M. Falcon, J. M. Gardiner, N. D. Miller, M. C. Romay, R. Walls, R. Walton, C.-T. Yeh, M. Bohn, et al. Maize genomes to fields: 2014 and 2015 field season genotype, phenotype, environment, and inbred ear image datasets. BMC Research Notes, 11(1):452, 2018.

[2] D. Alpern, V. Gardeux, J. Russeil, B. Mangeat, A. C. A. Meireles-Filho, R. Breysse, D. Hacker, and B. Deplancke. BRB-seq: ultra-affordable high-throughput transcriptomics enabled by bulk RNA bar-coding and sequencing. Genome Biology, 20:71, 2019. doi: 10.1186/s13059-019-1671-x.

[3] C. M. Andorf, J. Ross-Ibarra, A. S. Seetharam, M. B. Hufford, and M. R. Woodhouse. A unified VCF dataset from nearly 1,500 diverse maize accessions and resources to explore the genomic landscape of maize. G3: Genes, Genomes, Genetics, 15(2):jkae281, 2025. doi: 10.1093/g3journal/jkae281.

[4] C. B. Azodi, J. Pardo, R. VanBuren, G. de los Campos, and S.-H. Shiu. Transcriptome-based prediction of complex traits in maize. The Plant Cell, 32(1):139–151, 2020. doi: 10.1105/tpc.19.00332.

[5] N. L. Bray, H. Pimentel, P. Melsted, and L. Pachter. Near-optimal probabilistic RNA-seq quantification. Nature Biotechnology, 34(5):525–527, 2016. doi: 10.1038/nbt.3519.

[6] R. Bukowski, X. Guo, Y. Lu, C. Zou, B. He, Z. Rong, B. Wang, D. Xu, B. Yang, C. Xie, et al. Construction of the third-generation Zea mays haplotype map. GigaScience, 7(4):gix134, 2018. doi: 10.1093/gigascience/gix134.

[7] Q. Chen, J. D. Washburn, D. C. Lima, M. C. Romay, J. L. Gage, J. B. Holland, A. Xavier, S. C. Murray, D. Ertl, M. Lopez-Cruz, et al. Genomes to fields 2024 maize genotype by environment prediction competition. BMC Research Notes, 19(1):113, 2026. doi: 10.1186/s13104-026-07629-5.

[8] H. S. Chung, A. J. Koo, X. Gao, S. Jayanty, B. Thines, A. D. Jones, and G. A. Howe. Regulation and function of Arabidopsis JASMONATE ZIM-domain genes in response to wounding and herbivory. Plant Physiology, 146(3):952–964, 2008.

[9] V. Coustham, P. Li, A. Strange, C. Lister, J. Song, and C. Dean. Quantitative modulation of polycomb silencing underlies natural variation in vernalization. Science, 337(6094):584–587, 2012.

[10] M. Creach, B. Webster, L. Newton, J. Turkus, J. C. Schnable, A. Thompson, and R. VanBuren. Predicting complex phenotypes using multi-omics data in maize. The Plant Cell, 38(6):koag185, 2026. doi: 10.1093/plcell/koag185.

[11] P. Danecek, J. K. Bonfield, J. Liddle, J. Marshall, V. Ohan, M. O. Pollard, A. Whitwham, T. Keane, S. A. McCarthy, R. M. Davies, and H. Li. Twelve years of SAMtools and BCFtools. GigaScience, 10 (2):giab008, 2021. doi: 10.1093/gigascience/giab008.

[12] B. W. Diers, J. Specht, K. M. Rainey, P. Cregan, Q. Song, V. Ramasubramanian, G. Graef, R. Nelson, W. Schapaugh, D. Wang, G. Shannon, L. McHale, S. K. Kantartzi, A. Xavier, R. M. Mian, R. M. Stupar, J.-M. Michno, Y.-Q. C. An, W. Goettel, R. Ward, C. Fox, A. E. Lipka, D. Hyten, T. Cary, and W. D. Beavis. Genetic architecture of soybean yield and agronomic traits. G3: Genes, Genomes, Genetics, 8(10):3367–3375, 2018. doi: 10.1534/g3.118.200332.

[13] S. Griebel, A. Adedayo, and M. R. Tuinstra. Genetic diversity for starch quality and alkali spreading value in sorghum. The Plant Genome, 14(1):e20067, 2021. doi: 10.1002/tpg2.20067.

[14] M. W. Grzybowski and J. C. Schnable. Haplotype-rich cis-regulation underlies transcriptomic diversity across the breeding history of maize (Zea mays). bioRxiv, 2026. doi: 10.64898/2026.02.19.706772. Preprint.

[15] M. W. Grzybowski, R. V. Mural, G. Xu, J. Turkus, J. Yang, and J. C. Schnable. A common resequencing-based genetic marker data set for global maize diversity. The Plant Journal, 113(6): 1109–1121, 2023. doi: 10.1111/tpj.16123.

[16] M. Heyer, M. Reichelt, and A. Mithöfer. A holistic approach to analyze systemic jasmonate accumulation in individual leaves of Arabidopsis rosettes upon wounding. Frontiers in Plant Science, 9:1569, 2018.

[17] P. Y. Hsu and S. L. Harmer. Circadian phase has profound effects on differential expression analysis. PLOS ONE, 7(11):e49853, 2012.

[18] M. B. Hufford, A. S. Seetharam, M. R. Woodhouse, et al. De novo assembly, annotation, and comparative analysis of 26 diverse maize genomes. Science, 373(6555):655–662, 2021. doi: 10.1126/science.abg5289.

[19] M. Kashima, R. L. Sakamoto, H. Saito, S. Ohkubo, A. Tezuka, A. Deguchi, Y. Hashida, Y. Kurita, K. Iwayama, S. Adachi, et al. Genomic basis of transcriptome dynamics in rice under field conditions. Plant and Cell Physiology, 62(9):1436–1445, 2021.

[20] K. Kontolati, R. J. Gladstone, I. W. Davis, and E. M. Pickering. Biology-informed neural networks learn nonlinear representations from omics data to improve genomic prediction and biological discovery. The Plant Journal, 127(3):e71076, 2026.

[21] K. A. G. Kremling, S.-Y. Chen, M.-H. Su, N. K. Lepak, M. C. Romay, K. L. Swarts, F. Lu, A. Lorant, P. J. Bradbury, and E. S. Buckler. Dysregulation of expression correlates with rare-allele burden and fitness loss in maize. Nature, 555(7697):520–523, 2018. doi: 10.1038/nature25966.

[22] X. Lai, C. Bendix, L. Yan, Y. Zhang, J. C. Schnable, and F. G. Harmon. Interspecific analysis of diurnal gene regulation in panicoid grasses identifies known and novel regulatory motifs. BMC Genomics, 21 (1):428, 2020. doi: 10.1186/s12864-020-06824-3.

[23] D. Li, Q. Liu, and P. S. Schnable. TWAS results are complementary to and less affected by linkage disequilibrium than GWAS. Plant Physiology, 186(4):1800–1811, 2021. doi: 10.1093/plphys/kiab161.

[24] D. Li, Q. Wang, Y. Tian, X. Lyv, H. Zhang, H. Hong, H. Gao, Y.-F. Li, C. Zhao, J. Wang, et al. TWAS facilitates gene-scale trait genetic dissection through gene expression, structural variations, and alternative splicing in soybean. Plant Communications, 5(10):101010, 2024. doi: 10.1016/j.xplc.2024.101010.

[25] H. Li. Aligning sequence reads, clone sequences and assembly contigs with BWA-MEM. arXiv:1303.3997, 2013.

[26] Z. Liang, Z. A. Myers, D. Petrella, J. Engelhorn, T. Hartwig, and N. M. Springer. Mapping responsive genomic elements to heat stress in a maize diversity panel. Genome Biology, 23(1):234, 2022.

[27] D. C. Lima, J. D. Washburn, J. I. Varela, Q. Chen, J. L. Gage, M. C. Romay, J. B. Holland, D. Ertl, M. Lopez-Cruz, F. M. Aguate, G. de los Campos, S. M. Kaeppler, N. de León, et al. Genomes to Fields 2022 maize genotype by environment prediction competition. BMC Research Notes, 16(1):148, 2023. doi: 10.1186/s13104-023-06421-z.

[28] H.-J. Liu, X. Wang, Y. Xiao, J. Luo, F. Qiao, W. Yang, R. Zhang, Y. Meng, J. Sun, S. Yan, Y. Peng, L. Niu, L. Jian, W. Song, J. Yan, C. Li, Y. Zhao, Y. Liu, M. L. Warburton, J. Zhao, and J. Yan. CUBIC: an atlas of genetic architecture promises directed maize improvement. Genome Biology, 21(1):20, 2020. doi: 10.1186/s13059-020-1930-x.

[29] B. K. Lohman, J. N. Weber, and D. I. Bolnick. Evaluation of TagSeq, a reliable low-cost alternative for RNAseq. Molecular Ecology Resources, 16(6):1315–1321, 2016. doi: 10.1111/1755-0998.12529.

[30] M. I. Love, W. Huber, and S. Anders. Moderated estimation of fold change and dispersion for RNA-seq data with DESeq2. Genome Biology, 15(12):550, 2014. doi: 10.1186/s13059-014-0550-8.

[31] H. Mangal, K. Linders, J. Turkus, N. Shrestha, B. Long, E. Cebert, X. Kuang, J. V. Torres-Rodriguez, and J. C. Schnable. Genes and pathways determining flowering time variation in temperate-adapted sorghum. The Plant Journal, 122(5):e70250, 2025. doi: 10.1111/tpj.70250.

[32] M. Mazaheri, M. Heckwolf, B. Vaillancourt, J. L. Gage, B. Burdo, S. Heckwolf, K. Barry, A. Lipzen, C. B. Ribeiro, T. J. Y. Kono, H. F. Kaeppler, E. P. Spalding, C. N. Hirsch, C. R. Buell, N. de Leon, and S. M. Kaeppler. Genome-wide association analysis of stalk biomass and anatomical traits in maize. BMC Plant Biology, 19(1):45, 2019. doi: 10.1186/s12870-019-1653-x.

[33] G. Melandri, E. Monteverde, D. Riewe, H. AbdElgawad, S. R. McCouch, and H. Bouwmeester. Can biochemical traits bridge the gap between genomics and plant performance? a study in rice under drought. Plant Physiology, 189(2):1139–1152, 2022.

[34] E. J. Millet, W. Kruijer, A. Coupel-Ledru, S. Alvarez Prado, L. Cabrera-Bosquet, S. Lacube, A. Charcosset, C. Welcker, F. van Eeuwijk, and F. Tardieu. Genomic prediction of maize yield across European environmental conditions. Nature Genetics, 51(6):952–956, 2019. doi: 10.1038/s41588-019-0414-y.

[35] G. P. Morris, A. M. Harder, A. L. Healey, C. M. McLaughlin, J. L. Rifkin, C. Cruet-Burgos, J. W. Jenkins, S. Shu, J. J. Spiekerman, C. J. VanGessel, et al. A sorghum pangenome reference improves global crop trait discovery. Nature, 652(8112):1245–1253, 2026. doi: 10.1038/s41586-026-10229-9.

[36] A. J. Nagano, Y. Sato, M. Mihara, B. A. Antonio, R. Motoyama, H. Itoh, Y. Nagamura, and T. Izawa. Deciphering and prediction of transcriptome dynamics under fluctuating field conditions. Cell, 151(6): 1358–1369, 2012.

[37] R Core Team. R: A Language and Environment for Statistical Computing. R Foundation for Statistical Computing, Vienna, Austria, 2024. URL https://www.R-project.org/.

[38] P. Reymond, H. Weber, M. Damond, and E. E. Farmer. Differential gene expression in response to mechanical wounding and insect feeding in Arabidopsis. The Plant Cell, 12(5):707–719, 2000.

[39] C. Soneson, M. I. Love, and M. D. Robinson. Differential analyses for RNA-seq: transcript-level estimates improve gene-level inferences. F1000Research, 4:1521, 2015. doi: 10.12688/f1000research.7563.1.

[40] J. V. Torres-Rodríguez, D. Li, J. Turkus, L. Newton, J. Davis, L. Lopez-Corona, W. Ali, G. Sun, R. V. Mural, M. W. Grzybowski, B. M. Zamft, A. M. Thompson, and J. C. Schnable. Population-level gene expression can repeatedly link genes to functions in maize. The Plant Journal, 119(2):844–860, 2024. doi: 10.1111/tpj.16801.

