## Supplementary Information: Tables S1-S3, Figures S1-S3 for "Rapid, replicated field tissue collection for population-scale plant omics"

### Supplementary tables

Table S1: **RNA quality control statistics for samples employed for comparison of liquid nitrogen, dry ice, and RNA preservative collection methods.**

| | | Liquid nitrogen | | Dry ice | | RNA preservative | |
|---|---|---|---|---|---|---|---|
| Plot | Genotype | RNA (ng/$\mu$L) | RIN | RNA (ng/$\mu$L) | RIN | RNA (ng/$\mu$L) | RIN |
| 3501A | B14A/MO17 | 176.2 | 7.2 | 185.7 | 7.1 | – | – |
| 3502A | B73/MO17 | 119.2 | 6.6 | 179.9 | 7.0 | – | – |
| 3503A | CG119/CG108 | 109.2 | 6.1 | 91.7 | 6.0 | – | – |
| 3504A | REDEAR/LH195 | 203.8 | 6.9 | 232.3 | 6.9 | – | – |
| 3505A | F42/OH43 | 196.6 | 7.0 | 207.7 | 7.0 | – | – |
| 3506A | B73/TX779 | 246.8 | 7.1 | 240.1 | 6.9 | – | – |
| 3507A | CG44/CGR01 | 135.2 | 6.3 | 184.4 | 6.9 | – | – |
| 3508A | B14A/OH43 | 184.8 | 7.6 | 214.9 | 7.0 | – | – |
| 3509A | B37/OH43 | 163.4 | 7.3 | 166.8 | 7.5 | – | – |
| 3510A | PHW52/PHM49 | 189.7 | 7.1 | 188.5 | 7.7 | – | – |
| 3511A | PHW52/PHN82 | 216.6 | 7.1 | 221.0 | 7.5 | – | – |
| 3512A | PHG39/PHN82 | 219.4 | 8.1 | 176.2 | 7.5 | – | – |
| 3513A | LH74/PHN82 | 188.2 | 7.6 | 237.5 | 8.2 | – | – |
| 3514A | B37/H95 | 127.0 | 7.4 | 133.1 | 7.3 | – | – |
| 3515A | F42/H95 | 163.3 | 7.6 | 149.2 | 7.6 | – | – |
| 3516A | B73/PHN82 | 194.1 | 7.1 | 203.3 | 7.5 | – | – |
| 3517A | B73/TX779 | 150.8 | 7.4 | 161.1 | 7.1 | – | – |
| 3518A | B14A/H95 | 160.2 | 7.5 | 151.0 | 7.2 | – | – |
| 3519A | TX777/LH195 | 196.4 | 7.4 | 189.8 | 7.9 | – | – |
| 3520A | TX714/TX779 | 167.4 | 7.5 | 181.7 | 7.4 | – | – |
| 3521A | B73/PHM49 | 206.9 | 7.3 | 192.2 | 7.6 | – | – |
| 3522A | TX110/87916 | 95.2 | 6.8 | 115.1 | 7.2 | – | – |
| 3523A | F42/MO17 | 173.2 | 7.2 | 173.5 | 7.2 | – | – |
| 3524A | B37/MO17 | 144.9 | 7.1 | 170.0 | 7.2 | – | – |
| 3525A | 2369/LH123HT | 144.8 | 7.2 | 139.3 | 7.2 | – | – |
| Sample-1 | | 150.2 | 6.4 | 136.4 | 6.3 | 99.1 | 4.8 |
| Sample-2 | | 134.0 | 6.1 | 134.9 | 6.4 | 106.4 | 4.7 |
| Sample-3 | | 143.5 | 6.7 | 127.5 | 6.4 | 126.9 | 4.2 |

Table S2: **Detailed outcomes from between-plot carryover analysis.**

| | Genotype | | | Forward (%) | | | Reciprocal (%) | | | |
|---|---|---|---|---|---|---|---|---|---|---|
| Sampler | Sample 1 | Sample 2 | Markers | Ref | Alt | Mean | Ref | Alt | Mean | Third allele (%) |
| 1 | R101 | T146 | 20,842 | 0.1359 | 0.0365 | 0.0862 | 0.1015 | 0.0252 | 0.0634 | 0.0268 |
| 2 | R113 | W24 | 26,987 | 0.0843 | 0.0240 | 0.0541 | 0.0835 | 0.0228 | 0.0532 | 0.0263 |
| 3 | PHG83 | NC264 | 12,598 | 0.1813 | 0.0558 | 0.1185 | 0.2377 | 0.0921 | 0.1649 | 0.0315 |
| 5 | DK78004 | ND249 | 25,489 | 0.0959 | 0.0274 | 0.0616 | 0.1268 | 0.0333 | 0.0800 | 0.0343 |
| 6 | LH183 | L_155 | 27,969 | 0.1006 | 0.0394 | 0.0700 | 0.1147 | 0.0353 | 0.0750 | 0.0258 |
| 7 | PHJ65 | B87 | 14,851 | 0.1739 | 0.0658 | 0.1199 | 0.1016 | 0.0265 | 0.0641 | 0.0309 |
| 10 | 87916W | H49 | 14,611 | 0.1346 | 0.0496 | 0.0921 | 0.1118 | 0.0278 | 0.0698 | 0.0292 |
| Total / mean | | | 143,347 | 0.1295 | 0.0426 | 0.0861 | 0.1254 | 0.0376 | 0.0815 | 0.0293 |

Table S3: **Languages in which the written protocols, fabrication videos, and translations of this manuscript are available.**

| Language | Narrator | Fabrication protocol (Protocol S1) | Field collection protocol (Protocol S2) | Fabrication video | Translated paper | Native speakers (millions) |
|---|---|---|---|---|---|---|
| English | Jonathan Turkus | Protocol_S1_kit_fabrication_English.pdf | Protocol_S2_field_collection_English.pdf | YouTube link | — | 372 |
| Bengali | Pritha Kundu | Protocol_S1_kit_fabrication_Bengali.pdf | Protocol_S2_field_collection_Bengali.pdf | YouTube link | PDF link | 232 |
| Chinese | Zhongjie Ji | Protocol_S1_kit_fabrication_Chinese_Simplified.pdf | Protocol_S2_field_collection_Chinese_Simplified.pdf | YouTube link | PDF link | 988 |
| French | James Schnable (Voice Cloned) | Protocol_S1_kit_fabrication_French.pdf | Protocol_S2_field_collection_French.pdf | YouTube link | PDF link | 76 |
| German | Anna Podgórska-Grzybowska | Protocol_S1_kit_fabrication_German.pdf | Protocol_S2_field_collection_German.pdf | YouTube link | PDF link | 76 |
| Hindi | Harshita Mangal | Protocol_S1_kit_fabrication_Hindi.pdf | Protocol_S2_field_collection_Hindi.pdf | YouTube link | PDF link | 347 |
| Italian | James Schnable (Voice Cloned) | Protocol_S1_kit_fabrication_Italian.pdf | Protocol_S2_field_collection_Italian.pdf | YouTube link | PDF link | 60 |
| Japanese | Masako Obata | Protocol_S1_kit_fabrication_Japanese.pdf | Protocol_S2_field_collection_Japanese.pdf | YouTube link | PDF link | 124 |
| Kannada | Muragesh M. Hiremath | Protocol_S1_kit_fabrication_Kannada.pdf | Protocol_S2_field_collection_Kannada.pdf | YouTube link | PDF link | 44 |
| Korean | Fotima Turdalieva | Protocol_S1_kit_fabrication_Korean.pdf | Protocol_S2_field_collection_Korean.pdf | YouTube link | PDF link | 82 |
| Nepali | Bibash Rokaya | Protocol_S1_kit_fabrication_Nepali.pdf | Protocol_S2_field_collection_Nepali.pdf | YouTube link | PDF link | 19 |
| Persian | James Schnable (Voice Cloned) | Protocol_S1_kit_fabrication_Persian.pdf | Protocol_S2_field_collection_Persian.pdf | YouTube link | PDF link | 65 |
| Polish | Marcin Grzybowski | Protocol_S1_kit_fabrication_Polish.pdf | Protocol_S2_field_collection_Polish.pdf | YouTube link | PDF link | 40 |
| Portuguese | Henrique Moura Dias | Protocol_S1_kit_fabrication_Portuguese_Brazil.pdf | Protocol_S2_field_collection_Portuguese_Brazil.pdf | YouTube link | PDF link | 252 |
| Punjabi (Shahmukhi) | Waqar Ali | Protocol_S1_kit_fabrication_Punjabi_Shahmukhi.pdf | Protocol_S2_field_collection_Punjabi_Shahmukhi.pdf | YouTube link | PDF link | 90 |
| Rajasthani (Marwari) | Anuradha Choudhary | Protocol_S1_kit_fabrication_Rajasthani_Marwari.pdf | Protocol_S2_field_collection_Rajasthani_Marwari.pdf | YouTube link | PDF link | 21 |
| Russian | Fotima Turdalieva | Protocol_S1_kit_fabrication_Russian.pdf | Protocol_S2_field_collection_Russian.pdf | YouTube link | PDF link | 133 |
| Spanish | Karla Montserrat Cuéllar Pérez | Protocol_S1_kit_fabrication_Spanish.pdf | Protocol_S2_field_collection_Spanish.pdf | YouTube link | PDF link | 487 |
| Tamil | Surya Krishna Sakthivel | Protocol_S1_kit_fabrication_Tamil.pdf | Protocol_S2_field_collection_Tamil.pdf | YouTube link | PDF link | 79 |
| Telugu | Sai Subhash Mahamkali VS | Protocol_S1_kit_fabrication_Telugu.pdf | Protocol_S2_field_collection_Telugu.pdf | YouTube link | PDF link | 83 |
| Turkish | Özgür Altundaş | Protocol_S1_kit_fabrication_Turkish.pdf | Protocol_S2_field_collection_Turkish.pdf | YouTube link | PDF link | 86 |
| Urdu | Waqar Ali | Protocol_S1_kit_fabrication_Urdu.pdf | Protocol_S2_field_collection_Urdu.pdf | YouTube link | PDF link | 78 |
| Uzbek | Shokhista Turdalieva | Protocol_S1_kit_fabrication_Uzbek.pdf | Protocol_S2_field_collection_Uzbek.pdf | YouTube link | PDF link | 36 |
| Total | | | | | | 3,870 |

All protocol PDFs and design files are available from figshare (https://doi.org/10.6084/m9.figshare.35094379). All fabrication videos are collected in the YouTube playlist https://www.youtube.com/watch?v=Ozwe9-aWtT0&list=PLGAixDAU2zuQ.

### Supplementary figures

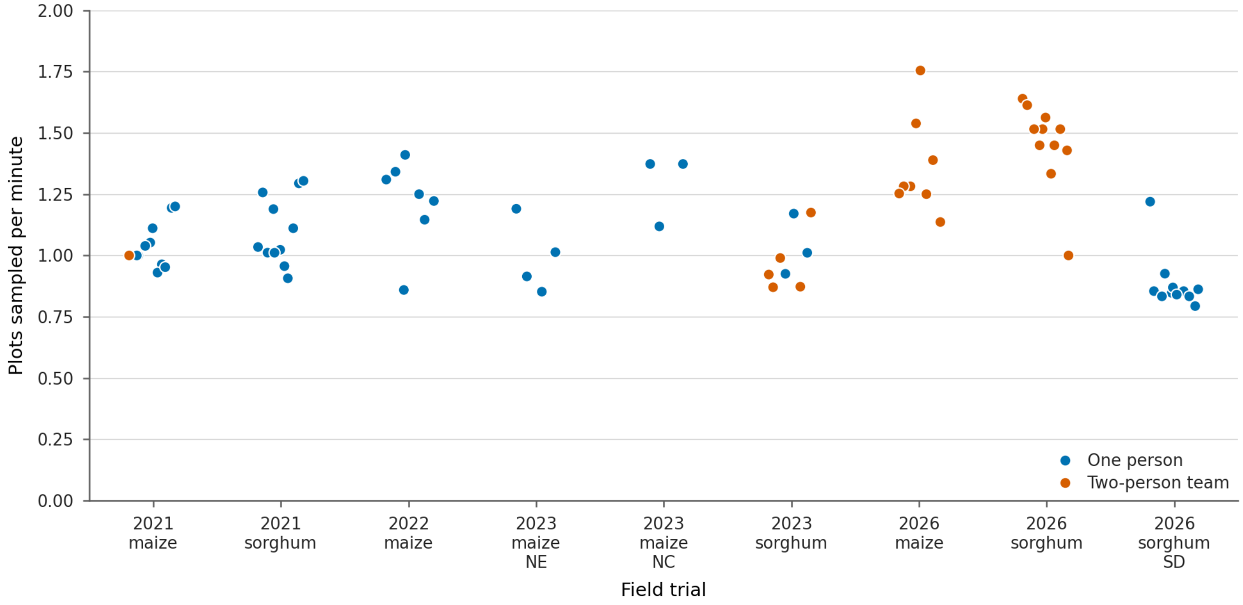


Figure S1: **Collection rates of individual samplers or teams across the nine field campaigns.** Each point represents either a single sampler or a two-person team using one collection kit.

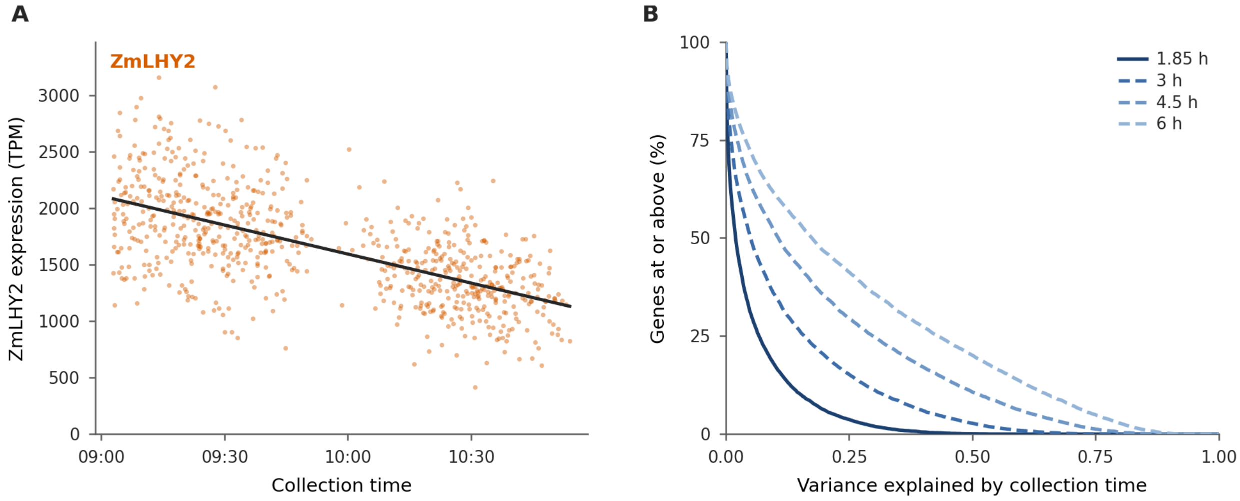


Figure S2: **Impact of collection time on the 2021 Nebraska maize dataset. (A)** Expression of *ZmLHY2* (Zm00001eb172450), one of two syntenic orthologs of *SbLHY* (Sobic.007G047400), in 743 RNA-seq libraries collected between 09:03 and 10:54. Unlike later campaigns, this collection did not have enough pre-loaded tube trays, so the gap reflects manual reloading between the first and second trays. **(B)** Cumulative share of 10,595 expressed genes (mean expression $\geq$ 10 TPM) for which collection time explains at least the proportion of variance shown on the x-axis. The solid line is the observed distribution over 1.85 h; dashed lines are linear projections across longer windows.

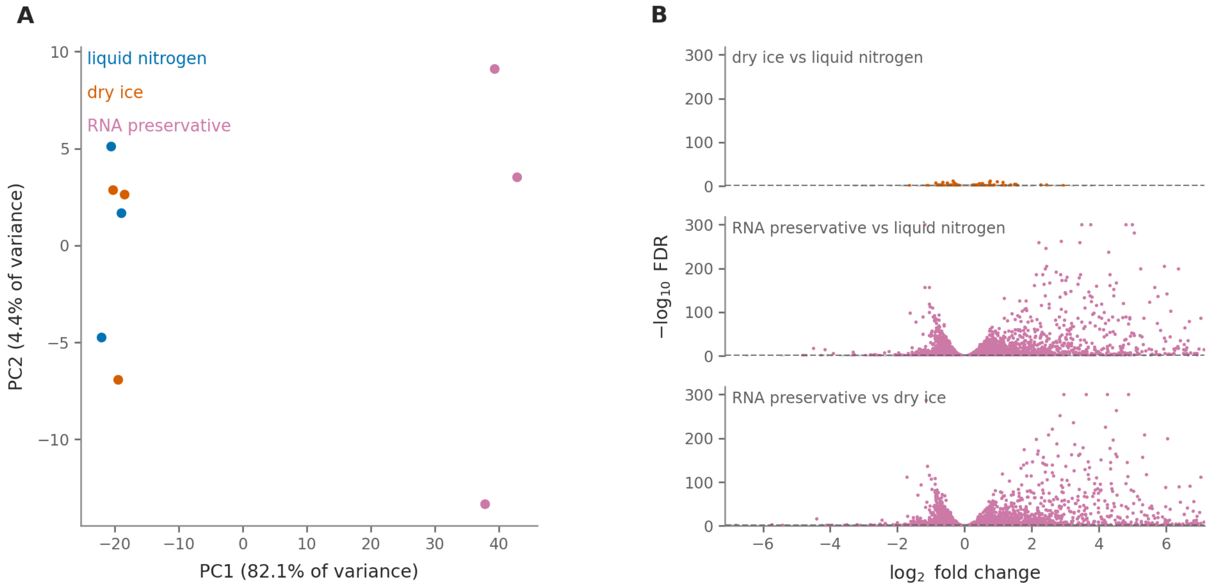


Figure S3: **Comparison of gene expression data from samples collected using different preservation strategies.** Results are from nine libraries, three each from liquid nitrogen, dry ice, and a commercial RNA preservative. **(A)** Principal-component analysis of the 500 most variable of 22,130 expressed genes on the $\log_2(\mathrm{TPM}+1)$ scale. **(B)** All three pairwise comparisons of differentially expressed genes using DESeq2 with Benjamini–Hochberg correction. A total of 138, 7,456, and 7,815 genes are differentially expressed at FDR $<0.05$ in the liquid nitrogen vs. dry ice, RNA preservative vs. liquid nitrogen, and RNA preservative vs. dry ice comparisons, respectively; 17, 1,108, and 1,157 exhibited at least a twofold change.

### Supplementary protocols

- **Supplementary Protocol S1:** Fabrication and assembly of the field tissue collection kit (triple leaf punch, cryogen carrier, tube tray, carrier box, and carry strap).
- **Supplementary Protocol S2:** Field collection procedure.
